# Enhanced prediction of protein functional identity through the integration of sequence and structural features

**DOI:** 10.1101/2024.09.30.615718

**Authors:** Suguru Fujita, Tohru Terada

## Abstract

Although over 300 million protein sequences are registered in a reference sequence database, only 0.2% have experimentally determined functions. This suggests that many valuable proteins, potentially catalyzing novel enzymatic reactions, remain undiscovered among the vast number of function-unknown proteins. In this study, we developed a method to predict whether two proteins catalyze the same enzymatic reaction by analyzing sequence and structural similarities, utilizing structural models predicted by AlphaFold2. We performed pocket detection and domain decomposition for each structural model. The similarity between protein pairs was assessed using features such as full-length sequence similarity, domain structural similarity, and pocket similarity. We developed several models using conventional machine learning algorithms and found that the LightGBM-based model outperformed the models. Our method also surpassed existing approaches, including those based solely on full-length sequence similarity and state-of-the-art deep learning models. Feature importance analysis revealed that domain sequence identity, calculated through structural alignment, had the greatest influence on the prediction. Therefore, our findings demonstrate that integrating sequence and structural information improves the accuracy of protein function prediction.

## 1. Introduction

Proteins fold into specific tertiary structures, enabling them to perform distinct biological functions. Among these, enzymes have increasingly been recognized as powerful tools in green chemistry due to their ability to catalyze chemical reactions under mild conditions, with high selectivity and reduced waste production [1]. As a result, significant efforts have been made to discover novel enzymes with valuable functions [2]. Now, let us consider the potential for finding such enzymes. The reference sequence (RefSeq) collection of the National Center for Biotechnology Information (NCBI) contains over 300 million protein sequences, yet the number of proteins with known functions is quite limited. For instance, the Swiss-Prot database, one of the most widely accessed sources of experimentally determined protein functions, includes only 570,000 entries, representing just 0.2% of known proteins. This indicates that a vast number of the proteins with unknown functions remain, and many useful enzymes are likely still undiscovered.

The functional information resistered in the Swiss-Prot database is sourced from scientific literatures and manually curated by experts. The protein’s function is described using Gene Ontology (GO) terms [4], Enzyme Comission (EC) numbers [5], Kyoto Encyclopedia of Genes and Genomes (KEGG) identifiers (IDs) [6], as well as through descriptive text. Enzymatic reactions catalyzed by the protein are associated with the entries of the Rhea database [7], which classifes enzymatic reactions by their reaction participants (i.e., their substrates and products) and assigns a unique ID (Rhea ID) to each class of the reaction.

Experimental determination of protein function is both expensive and time-consuming, which limits the number of proteins with known functions. To address this issue, researchers have developed computational methods to predict protein function from amino acid sequences [3]. Sequence similarity search programs, such as BLAST, are commonly used to predict protein function by comparing it to proteins with known functions [4,5]. However, these programs are less effective at predicting the function of enzymes, particularly their substrates and products, because small changes in amino acid sequences can affect substrate specificity.

Since an enzymes’s substrate specificity is determined by the spatial arrangement of amino acid residues in its substrate-binding pocket, incorporating structural information should enhance function prediction accuracy. Although structural information was limited to experimentally determined structures, typically available only for proteins with known functions. However, the advent of the deep learning-based protein structure prediction program AlphaFold2 (AF2) [6] has eliminated this limitation. AF2 has already predicted the structures of over 200 million proteins, now accessible through the AlphaFold Protein Structure Database (AFDB) [7]. Consequently, accurate enzyme function prediction is now acheivable using these predicted structures.

Deep learning techniques had also been directly applied to protein function prediction [8]. Various models have been developed to predict GO terms [9], EC classes [10–12], and functional domains within protein sequences [13]. In line with recent advances in deep learning, more complex models trained on large datasets have been used to improve protein function prediction methods. For example, the protein language model Evolutionary Scale Modeling (ESM), trained on millions of protein sequences, has demonstrated the ability to predict both the structure and function of proteins from their amino acid sequences [14,15]. Graph neural network-based models, such as DeepFRI [16] and GearNet [17], trained on extensive sequence and structural data, have also successfully predicted protein functions [18].

Although these models have achieved high prediction performance, they are difficult to interpret [19] and computationally expensive [20] due to their complexity. In contrast, conventional machine learning models with hypothesis-driven feature engineering provide better interpretability and require less computational power. Therefore, we adopted conventional machine learning models to develope protein function prediction methods. In this study, we constructed several binary classifiers to predict whether two proteins catalyze the same enzymatic reaction, training them using selected features representing sequence and structural similarities between the proteins. The prediction performances of these classifiers were compared with each other, with those based solely on sequence similarity, and with state-of-the-art deep learning-based methods. Additionally, we conducted feature importance analyses to assess the contribution of structural information to protein function prediction.

## 2. Materials and methods

### 2.1. Dataset construction

We obtained the sequences and structural models of 225,323 proteins associated with Rhea IDs, from Swiss-Prot (Release 2022_04) and AFDB (v4), respectively. Each protein sequence was subjected to a BLAST [26] search against the Swiss-Prot protein sequence database. Hits with E-values less than 10 that matched the 225,323 proteins were extracted to create a list of protein pairs. The protein pairs were then classified into two sets: one composed of protein pairs sharing identical functions, indicated by the same Rhea IDs, and the other composed of protein pairs having different functions with distinct RheaID. We constucted a dataset by randomly selecting 50,000 pairs with identical functions and 50,000 pairs with different functions. For each protein in the dataset, surface pockets were detected using Ghecom [21–23]. Pocket-forming residues were defined as those with non-hydrogen atoms located within 5 Å of a probe in a pocket, and these residues were extracted from each pocket. Subsequently, we decomposed the structural models into domains using the pae_to_domains program, which performs domain decomposition of AF2-predicted structures based on predicted aligned error (PAE) [24], employing a graph-based community clustering approach [25]. Only residues with per-residue confidence (pLDDT) greater than 70 were cosidered, and small domains with fewer than 30 residues were discarded.

### 2.2. Feature engineering

2.2.1. Full-length sequence similarity features The E-value and sequence identity calculated with BLAST [26] in the previous section were used as full-length sequence similarity features for each protein pair, referred to as Full_Evalue and Full_SeqID, respectively. The E-value was converted to its common logarithm, log_10_(E-value), if E-value > 10^−180^; otherwise, log_10_(E-value) was set to −200.

#### 2.2.2 Domain structural similarity features

Since AF2 cannot accurately predict the relative positions of structurally independent domains, we decomposed the structural models into domains based on PAE and assessed the structural similarity between protein pairs by comparing their domains. If proteins A and B, being compared, consist of *n*_A_ and *n*_B_ domains, respectively, structure alignment was performed using TM-align [27] for each of *n*_A_ × *n*_B_ pairs of the domains. For each domain pair, we calculated the average TM-score [28], root-mean-square deviation (RMSD), and sequence identity. The values from the domain pair with the highest average TM-score were selected as the structural similarity features for the protein pair. These features are referred to as Dmn_TM, Dmn_RMSD, and Dmn_SeqID, respectively.

#### 2.2.3. Pocket similarity features

A pocket within a domain was defined as the intersection of the pocket-forming residues and the residues comprising the domain. The similarity between pockets in two proteins was evaluated by comparing the pockets within the domains that had the highest average TM-score for the protein pair. If a domain contained two or more pockets, only the largest pocket was considered. Structure comparison was performed using PyMol’s “align” function, and the RMSD and sequence identity for the pocket pair were calculated. These values were used as the pocket similarity features for the protein pair, referred to as Pckt_RMSD and Pckt_SeqID, respectively.

### 2.3. Model training and hyperparameter optimization

After excluding protein pairs for which some similarity features could not be computed, the final dataset consisted of 20,800 pairs with identical functions and 20,800 pairs with different functions. The dataset was then randomly divided into training (56.25%), validation (18.75%), and test sub-datasets (25%), with each sub-dataset containing an equal number of identical-function and different-function protein pairs. We developed binary classification models using the LightGBM [29], Random Forest, Decision Tree, AdaBoost, Logistic Regression, Linear Discriminant Analysis, K-Nearest Neighbors, Naïve Bayes, and Support-Vector Machine algorithms to predict the functional identity between a pair of proteins, i.e., whether the two proteins share the same function. Each model was trained on the training sub-dataset, and its hyper-parameters were tuned using Optuna [31], an automatic hyperparameter search framework, through 5-fold cross validation on the validation sub-dataset. The area under the receiver operating characteristic curve (AUROC) was used as the evaluation metric during validation. To assess the stability and variability of the model, we conducted 50 bootstrap iterations.

### 2.4. Performance assessment

We used six common metrics to assess performance: accuracy (ACC), false positive rate (FPR), Matthews correlation coefficient (MCC), precision (PRE), recall (REC), and F1 score. The prediction threshold was set to maximize the F1 score. The metrics are defined as follows:

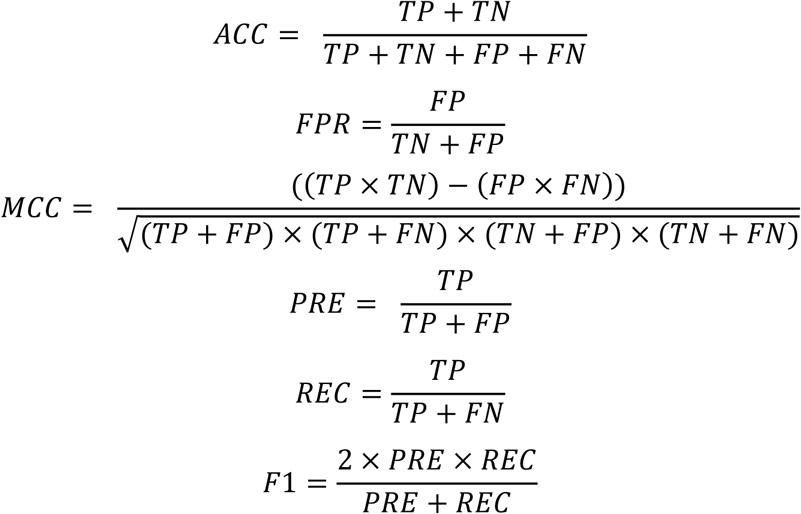

where *TP*, *FP*, *TN*, and *FN* represent the numbers of true positives, false positives, true negatives, and false negatives, respectively. We used the AUROC and the area under the precision-recall curve (AUPR) to compare the performance of different models.

### 2.5. Prediction with DeepFRI

DeepFRI uses graph convolutional neural networks to predict protein function based on sequence and structural information [31]. The model is pre-trained to predict both GO terms and EC numbers for proteins. DeepFRI generates a vector of real numbers between 0 and 1, where each number corresponds to a specific protein function (i.e., a GO term or EC number) and represents the prediction confidence that the input protein exhibits the corresponding function. In other words, DeepFRI predicts multiple functions for a protein simultaneously, along with their associated confidence levels. To compare the prediction performance of DeepFRI with our models, we calculated the cosine similarity between the output vectors for protein pairs, as described in [32]. This similarity measure was used to assess the plausibility that the two proteins have the same function. The pre-trained model was obtained from https://github.com/flatironinstitute/DeepFRI, and EC-number predictions were conducted using the PDB files of the model structures as input.

### 2.6. Prediction with ESM-2

The ESM-2 model [33] is a transformer-based protein language model [34] designed to generate numerical sequences from a protein’s amino acid sequence. It is pre-trained on the UniRef50 dataset, which comprises 3 million representative protein sequences [35]. The model outputs the hidden states of an input amino acid sequence of length *n* as a sequence of *d*-dimensional vectors (ℎ_1_, ⋯, ℎ_*n*_) (*d* = 1,280). The protein’s single representation is obtained by averaging these vectors, as follows:

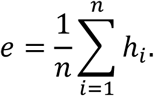

The likelihood that a pair of proteins share the same function was assessed by calculating the cosine similarity between their averaged vectors. This approach is based on the premise that proteins with similar sequences are likely to have similar functions. The pre-trained model used for this analysis was obtained from https://github.com/facebookresearch/esm (version esm2_t33_650M_UR50D).

### 2.7. Feature importance

#### 2.7.1. Gini importance

We calculated the Gini importance of each feature to assess its effectiveness in classification for our best model using the training sub-dataset [36].

#### 2.7.2. Shapley additive explanations

We performed a Shapley additive explanations (SHAP) analysis for our best model [37]. This involved calculating the Shapley value for each feature of every protein pair in the test sub-dataset to evaluate its contribution to the output value. The importance of each feature was determined by averaging the absolute Shapley value across the dataset.

## 3. Results and discussion

### 3.1 Data set construction and feature calculation

To explore the relationship between sequence similarity and the probability of having identical functions, we analyzed the result from the initial BLAST searches conducted for each of the 225,323 full-length protein sequences with Rhea IDs against the Swiss-Prot protein sequence database. The plot showing the rate of protein pairs with identical functions as a function of their log_10_(E-value) reveals that more than 50% of protein pairs have identical functions when the E-value is less than 10^−10^ (Fig. 1). This finding supports a binary classification approach where if the E-value between a pair of proteins is below a certain threshold, the proteins are predicted to have identical functions. This prediction method is known as the E-value-based method. We assessed the performance of this method and compared it with the method described below.

**Fig. 1.**
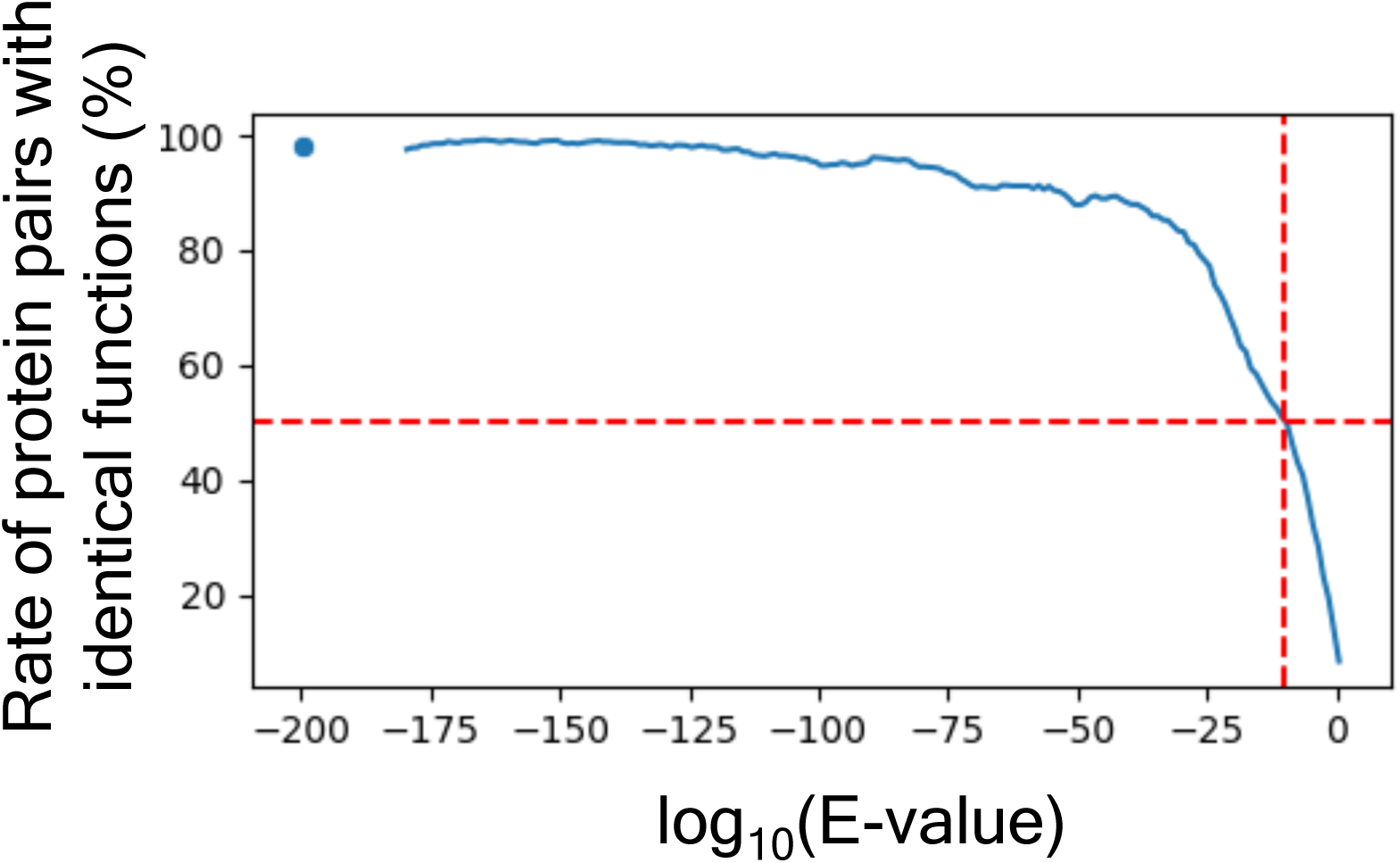
Plot of the rate of protein pairs with identical functions as a function of their log_10_(E-value). The red dashed lines indicates that the rate is 50% at log_10_(E-value) = −10.

To achieve better prediction performance, we developed machine learning models that utilize both sequence and structural information to predict the functional identity of protein pairs. We constructed a dataset comprising 20,800 protein pairs with identical functions and 20,800 protein pairs with different functions by selecting protein pairs with significant (E-value < 10) sequence similarities. It is important to note that even protein pairs with different functions can exhibit significant sequence similarities. Consequently, our prediction method does not target protein pairs with no sequence similarities, as these are typically functionally distinct. Each protein pair in the dataset was characterized by two full-length sequence similarity features (Full_Evalue and Full_SeqID), three domain structural similarity features (Dmn_TM, Dmn_RMSD, and Dmn_SeqID), and two pocket similarity features (Pckt_RMSD and Pckt_SeqID) (Table S1). We compared the distributions of these feature values between protein pairs with identical functions and those with different functions (Fig. S1). Overall, protein pairs with identical functions exhibited better similarity scores (i.e., smaller Full_Evalue, Dmn_RMSD, and Pckt_RMSD, and larger Full_SeqID, Dmn_TM, Dmn_SeqID, and Pckt_SeqID) compared to pairs with different functions. This suggests that models trained with these features could enhance the accuracy of functional identity predictions.

### 3.2 Model selection

We developed machine learning models using the LightGBM, Random Forest, Decision Tree, AdaBoost, Logistic Regression, Linear Discriminant Analysis, K-Nearest Neighbors, Naïve Bayes, and Support-Vector Machine algorithms, and trained them with the training sub-dataset. We employed Optuna to fine-tune the hyperparameters of each model using 5-fold cross validation on the validation sub-dataset to maximize the AUROC value [36]. We then compared the performance of these optimized models on the test sub-dataset (Table S2). The LightGBM-based model achieved the highest AUROC value, making it the best-performing model and the one selected for further analysis.

### 3.3 Comparison with existing methods

We compared the prediction performance of our LightGBM-based method with existing methods. Figures 2A and 2B display the ROC and PR curves of our LightGBM-based method, the E-value-based method, the ESM-2-based method, and the DeepFRI-based method.

**Fig. 2.**
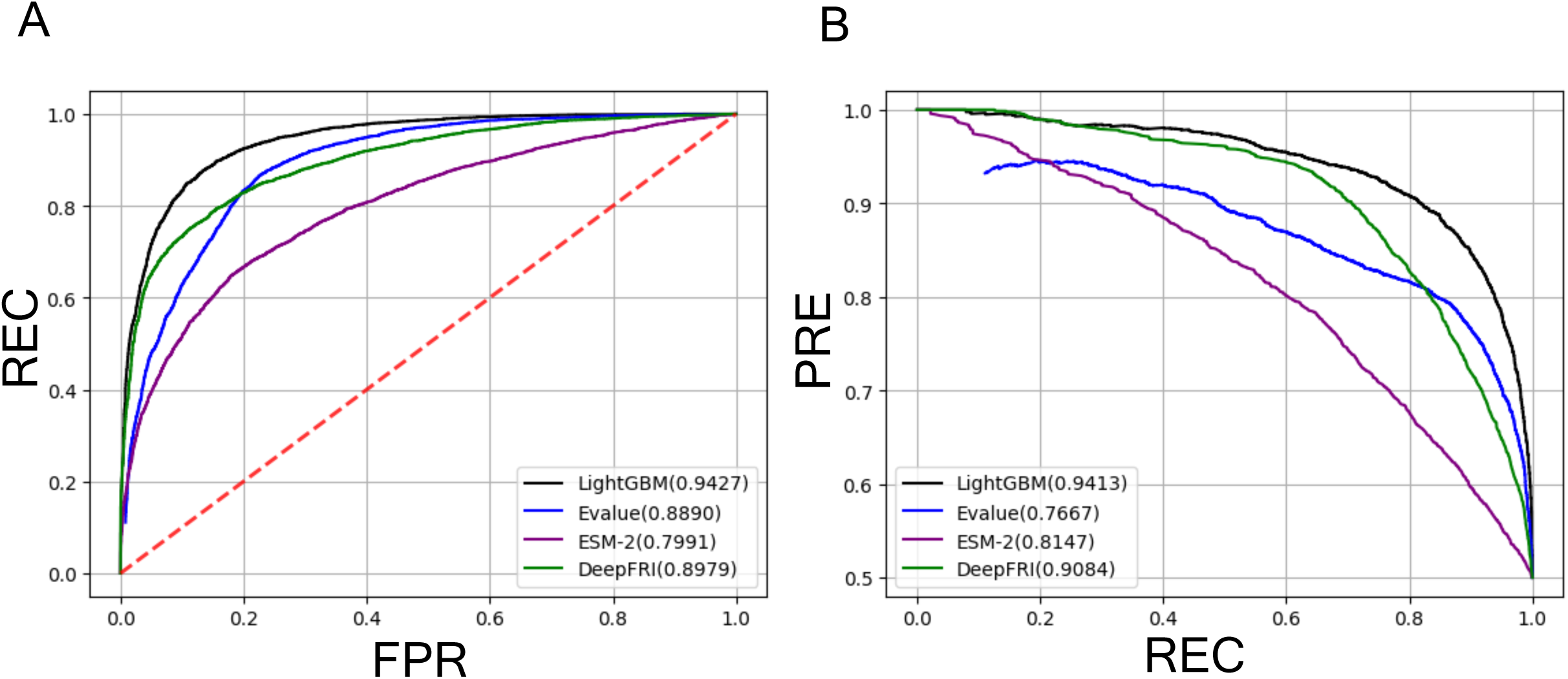
Model performance on the test sub-dataset. (A) Comparison of ROC curves for different methods, with the AUROC value for each method indicated in parenthesis. (B) Comparison of PR curves for different methods, with the AUPR value for each method indicated in parenthesis.

Additional performance metrics are summarized in Table 1. The ROC curve of our method outperformed those of the existing methods, indicating superior performance. Consistently, the AUROC value of our method (0.9427) exceeded those of the other methods (E-value: 0.8890; DeepFRI: 0.8963; ESM-2: 0.7991) (Table 1). Similarly, the PR curve of our method also surpassed the others, yielding a higher AURP value (our method: 0.9413) compared to the other methods (E-value: 0.7767; DeepFRI: 0.9067; ESM-2: 0.8147) (Table 1). Our model also showed better results in terms of ACC, FPR, MCC, PRE, REC, and F1 scores than the other models, except for REC, where the E-value-based method (87.75%) slightly outperformed our method (87.05%) (Table 1). The ROC plot curve indicates a relationship between REC and FPR. The slightly better REC value of the E-value-based method came at the expense of a much worse FPR (23.92%) compared to our method (13.35%). Additionally, we evaluated functional identity prediction using the Pfam database [38] and compared its performance with our method (Table 1). Similar to the E-value-based method, Pfam showed a better REC but a much worse FPR (Table 1), highlighting that enzymes within the same family or superfamily do not always target the same substrate. Overall, our method demonstrated superior prediction performance compared to the existing methods.

**Table 1.** Comparison of prediction performance among LightGBM and existing methods.

| Methods | ACC<br>(%) | PRE<br>(%) | REC<br>(%) | FPR (%) | F1 | MCC | AUROC | AUPR |
| --- | --- | --- | --- | --- | --- | --- | --- | --- |
| LightGBM | 87.05 | 87.24 | 87.05 | 13.35 | 0.8728 | 0.7421 | 0.9427 | 0.9413 |
| E-value | 81.91 | 78.58 | 87.75 | 23.92 | 0.8291 | 0.6427 | 0.8890 | 0.7767 |
| DeepFRI | 80.88 | 79.77 | 83.16 | 21.45 | 0.8143 | 0.6790 | 0.8963 | 0.9067 |
| ESM2 | 71.33 | 68.39 | 79.33 | 36.67 | 0.7345 | 0.4321 | 0.7991 | 0.8147 |
| Pfam | 61.99 | 56.92 | 98.60 | 74.62 | 0.7217 | 0.3520 | — | — |

### 3.4 Analysis of feature importance

As previously mentioned, conventional machine learning models, including LightGBM, offer greater model interpretability compared to deep learning-based models. To assess the effectiveness of each feature in our model for classification, we calculated the Gini importance (Fig. 3A). Our analysis revealed that Dmn_SeqID was the most influential feature, followed by Full_Evalue, Full_SeqID, Dmn_RMSD, and Dmn_TM. We next performed the SHAP analysis to evaluate the impact of each feature on the prediction result (Fig. 3B) [37]. We found that Dmn_SeqID had the greatest average impact on predictions. Consistently, the Shapley values for Dmn_SeqID used in the SHAP analysis were better correlated with its feature values (i.e., Dmn_SeqID) than those for Full_Evalue (Fig. S2). Since Dmn_SeqID is derived from the structural alignment of domains, these results underscore the importance of structural similarity in predicting functional identity.

**Fig. 3.**
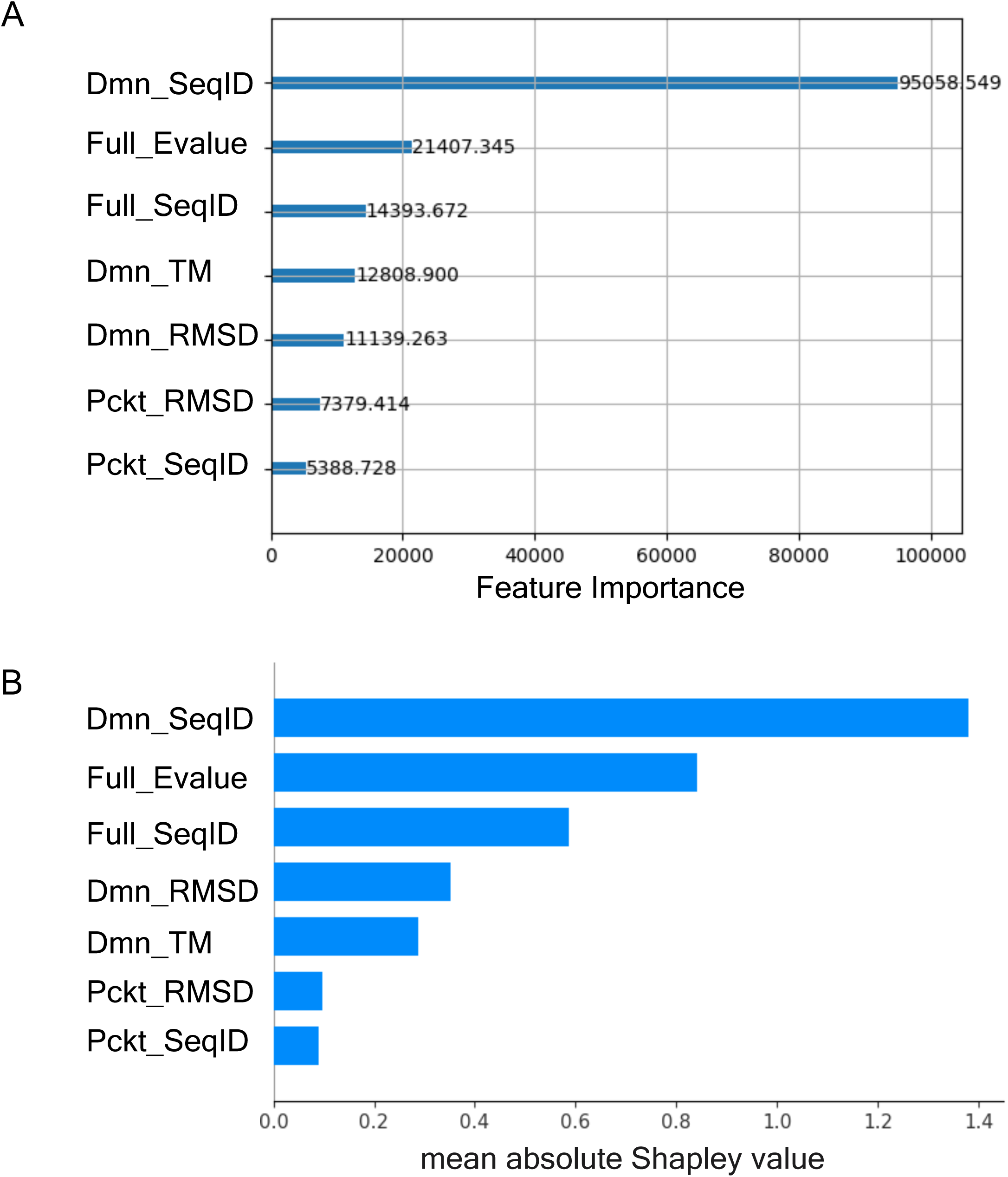
Comparisons of Gini importance calculated for the training sub-dataset (A) and mean absolute Shapley values calculated for the test sub-dataset (B) between features.

### 3.5 Performance on low sequence similarity dataset

We found that the Full_Evalue feature had the second highest Gini and SHAP importances (Fig. 3), indicating the significant role of full-length sequence similarity in function prediction. However, as noted earlier, the likelihood of identical functions drops below 50% when the E-value of full-length sequences of a pair of proteins exceeds 10^−10^ (Fig. 1). This suggests that relying solely on E-values for pairs with values larger than 10^−10^ makes it challenging to identify protein pairs with identical functions. To assess the performance of our method on such protein pairs, we created a new dataset and performed predictions on it. This new dataset, termed the low sequence similarity (LSS) dataset, includes protein pairs with E-values greater than 10^−10^ (comprising 680 pairs with identical functions and 680 pairs with different functions) and excludes protein pairs from the original dataset. Figures 4A and 4B present the ROC and PR curves of our method, the E-value-based method, the ESM-2-based method, and the DeepFRI-based method. As expected, the E-value-based method performed worse on the LSS dataset compared to the original test sub-dataset. The AUROC and AUPR values for the E-value-based method on the LSS dataset were 0.7391 and 0.7101, which were lower than those of our model (AUROC: 0.8701; AUPR: 0.8793). Similarly, the AUROC and AUPR values for the ESM-2-based and DeepFRI-based methods were also lower than those of our model (AUROC of DeepFRI: 0.8205; ESM-2: 0.5225, AUPR of DeepFRI: 0.8322; ESM-2: 0.5362).

**Fig. 4.**
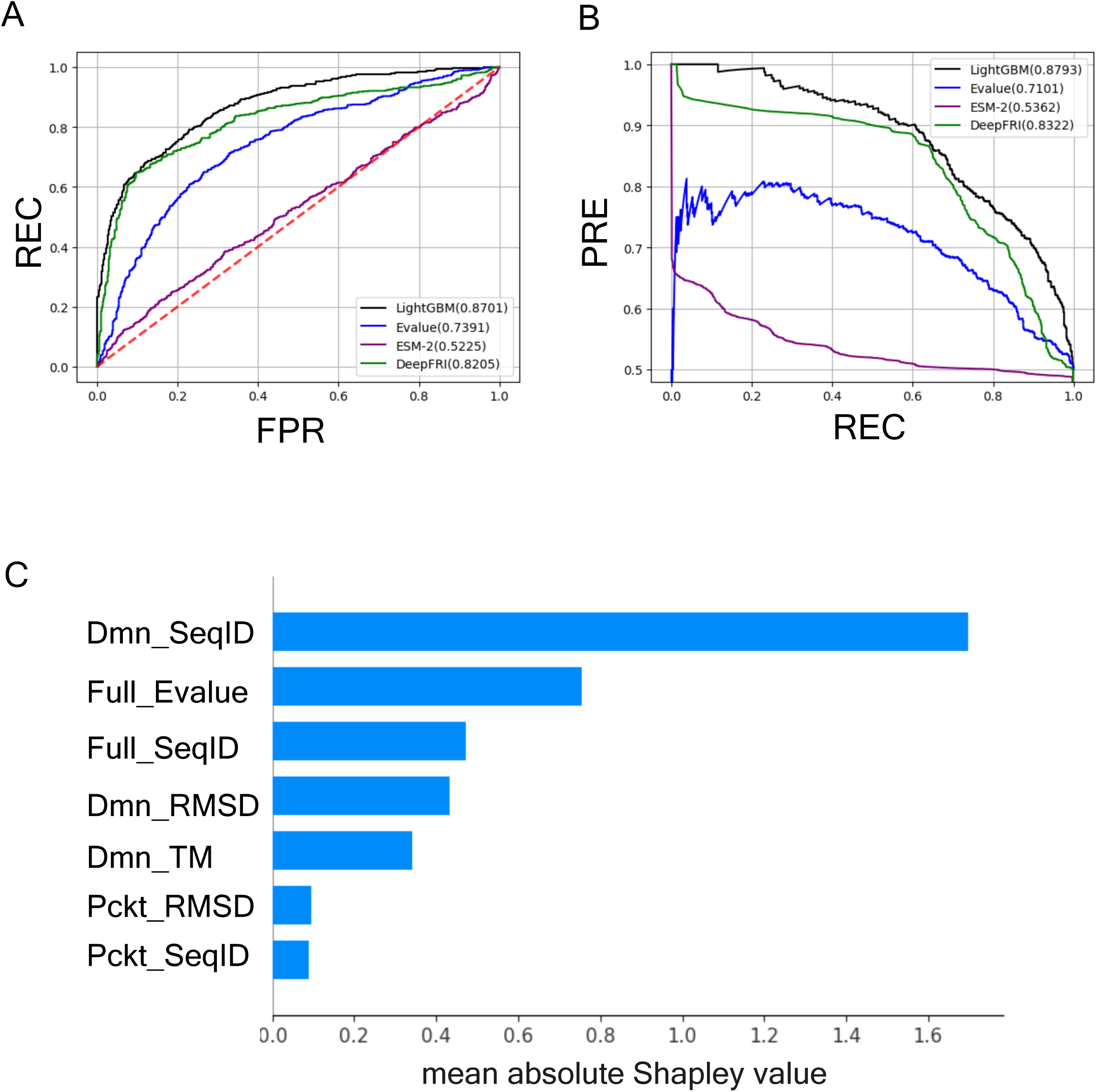
Model performance on the LSS dataset. (A) Comparison of ROC curves for different methods on the LSS dataset. (B) Comparison of PR curves for different methods on the LSS dataset. (C) Comparison of mean absolute Shapley values for features on the LSS dataset.

As with the original test sub-dataset, Dmn_SeqID had the largest average impact on predictions for the LSS dataset (Fig. 4C). These results demonstrate the robustness of our method against protein pairs with low full-length similarity and underscore the importance of structural similarity in function prediction.

### 3.6 Limitations and perspectives

In our method, the domains with the highest TM-score are selected from each protein pair to evaluate structural similarity. Consequently, if two proteins are predicted to have identical functions, this result does not mean that the domains chosen for structural comparison possess enzymatic activity. Other domains within the proteins may be responsible for the function. To address this issue, we plan to extend our method to include structural similarities among multiple domain pairs within a protein pair.

To enhance the practical application of our method, we are currently developing a database that will store sequences, domain structures, and pocket-forming residues of all function-known proteins. This database will enable us to predict the function of a protein by identifying other proteins with potentially identical functions. The output score of our method, which represents the confidence of the prediction, will allow us to rank hits by their confidence levels. Importantly, our method can also be applied to pairs of function-unknown proteins to predict their functional identities. Using our method, we can classify function-unknown proteins into clusters based on functional similarity, with the output score serving as an indicator of plausibility. This approach will help reduce the number of proteins requiring experimental functional determinations and contribute to elucidating the functions of previously uncharacterized proteins.

## 4. Conclusions

In this study, we developed a LightGBM-based method to predict the functional identity of protein pairs by utilizing full-length sequence similarity as well as the structural similarities of domains and pocket-forming residues between the proteins. Our method demonstrated superior prediction performance compared to methods based solely on full-length sequence similarity and those using ESM-2 and DeepFRI. LightGBM offers an advantage in model interpretability over deep learning models such as ESM-2 and DeepFRI. Feature importance analyses, including Gini importance and SHAP analysis, revealed that domain sequence identity, derived from structural alignment, had the most significant impact on predictions. These findings highlight the crucial role of structural similarity in predicting protein function.

### CRediT authorship contribution statement

**Suguru Fujita**: Conceptualization, Methodology, Software, Validation, Formal analysis, Investigation, Data curation, Visualization, Writing - original draft. **Tohru Terada**: Conceptualization, Methodology, Formal analysis, Investigation, Resources, Writing - review & editing, Supervision, Project administration, Funding acquisition.

### Declaration of Competing Interests

The authors declare that they have no known competing financial interests or personal relationships that could have perceived to influence the work reported in this paper.

## Acknowledgments

This work was partially supported by the JSPS KAKENHI Grant Number JP22H05126 and by Research Support Project for Life Science and Drug Discovery (Basis for Supporting Innovative Drug Discovery and Life Science Research (BINDS)) from Japan Agency for Medical Research and Development (AMED), under Grant Number JP24ama121027. The authors would like to thank Enago (www.enago.jp) for the English language review.

## Notes

### Competing Interest Statement

The authors have declared no competing interest.

